# Donor Sex and Platelet Storage Influence the Therapeutic Effects of Platelet-Derived Extracellular Vesicles on Endothelial Barrier Function

**DOI:** 10.1101/2025.10.23.683877

**Authors:** Mandeep Kaur, Malvika Gupta, Sowmya Shree Gopal, Charles E. Wade, Jessica C. Cardenas, Amit K. Srivastava

## Abstract

Platelet-derived extracellular vesicles (PEVs) play an active role in vascular protection and repair and are being explored as a viable alternative to platelet therapy. Because platelet function and stability are shaped by donor sex and storage conditions, these same factors are likely to influence the PEVs they release. Understanding these influences is key to developing PEVs into a safe and dependable therapeutic option. In this study, we investigated how donor sex and platelet storage affect the therapeutic properties of PEVs. To address this, PEVs were isolated from platelets of healthy male and female donors. Platelets were either processed immediately after blood collection to represent a resting state or stored overnight at room temperature on a rocker to mimic platelet storage conditions. PEVs isolated from these preparations displayed similar size, morphology, and cellular uptake across groups, but their biological effects diverged. Female PEVs, particularly from resting platelets, provided the strongest protection against thrombin-induced endothelial barrier disruption, stabilized junctional proteins, and reduced oxidative stress. Male PEVs showed weaker barrier protection compared to female-derived PEVs but more pronounced modulation of certain inflammatory mediators. In addition, PEVs derived from resting platelets (RP-PEVs) consistently showed stronger protective effects than those from stored platelets (SP-PEVs), regardless of donor sex. These results highlight that donor sex and platelet storage influence PEVs function and underscore the need to account for both when developing PEV-based therapies.

**Key Point:** The endothelial-protective effects of platelet-derived extracellular vesicles are modulated by platelet storage conditions and donor sex.

## 1. Introduction

Endothelial dysfunction, defined by structural and functional disruption of the vascular lining^1^, is a critical driver of vascular pathology^2^. Triggered by hypoxia, inflammation, and damage-associated molecular pattern (DAMP) molecules, it initiates a cascade of events including increased vascular permeability, dysregulated inflammatory signaling, and coagulation disturbances^3^. These processes underlie the progression of hemorrhagic shock, multi-organ dysfunction, and impaired tissue repair^4–7^. Injured endothelial cells further amplify oxidative stress, creating a self-perpetuating cycle of vascular damage^8^. Persistent endothelial barrier failure and systemic inflammation lead to adverse clinical outcomes^2,9–11^.

Preservation of endothelial integrity is essential for vascular homeostasis, and platelets are central to endothelial repair, promoting proliferation, stabilizing intercellular junctions, and maintaining barrier function^12,13^. Through the secretion of bioactive factors, platelets offer a unique therapeutic avenue for endothelial dysfunction^14^. However, the clinical utility of platelets therapy is limited by short shelf life (5 days), storage-related functional decline, and risks such as transfusion-related acute lung injury and alloimmune reactions^15^. These limitations highlight the need for alternative strategies that harness platelet therapeutic properties while overcoming logistical and safety concerns^16^.

Our group and others have demonstrated that platelet-derived extracellular vesicles (PEVs) preserve many of the biological properties of platelets and represent an alternative therapeutic strategy^4,16,17^. PEVs carry hemostatic proteins, transfer platelet receptors that activate signaling pathways^18^, and interact with endothelial cells and monocytes to modulate inflammatory responses^19^. They also deliver growth factors and microRNAs that promote endothelial proliferation, migration, and survival while reducing apoptosis^20–23^. Importantly, PEVs can be stored at low temperatures with preserved stability, addressing major limitations associated with platelet storage, transport, and shelf life^16,24^. These attributes position PEVs as viable substitutes for platelet transfusion, particularly in resource-limited or austere environments. PEVs activity, however, may be influenced by donor biology. For example, platelets from older males exhibit distinct metabolic profiles compared with those from younger males and females^25^, potentially shaping PEV composition and function. In addition, platelet storage is known to alter platelet functionality^26^, suggesting that both donor characteristics and storage conditions could impact PEVs efficacy. Our prior work showed that PEVs can restore endothelial integrity after vascular injury^4^; however, these findings were derived from a single donor, limiting generalizability.

In the present study, we evaluated the endothelial protective effects of PEVs obtained from multiple donors of both sexes and from platelets that were either freshly isolated or stored. Resting platelet-derived PEVs (RP-PEVs) were collected from platelet-rich plasma (PRP) immediately after blood collection, whereas stored platelet-derived PEVs (SP-PEVs) were obtained after PRP was maintained on a rocker for 24 hours at room temperature to mimic platelet storage conditions. We hypothesized that donor sex and storage condition influence the therapeutic properties of PEVs in regulating endothelial barrier function. To test this, we compared RP-PEVs and SP-PEVs from male and female donors to determine how these variables shape their vascular protective potential.

## 2. Materials and Methods

### 2.1. Donor recruitment, eligibility screening, and platelet collection procedures

Platelets were obtained from 20 healthy young adult donors (10 males and 10 females). We excluded subjects who were pregnant, had autoimmune or inflammatory disorders, or had used antiplatelet agents within the 14 days prior to blood collection. All donors were asked to abstain from alcohol and caffeine for 12 hours before blood collection. Venous blood was drawn using a 21-gauge needle with a free-flow method to help minimize platelet activation. Blood samples were placed into tubes containing acidic citrate dextrose (ACD) at a ratio of 1:8. PRP was then isolated as previously described^16,27^. The study was approved by the Institutional Review Board (IRB) of Thomas Jefferson University, Philadelphia, and written informed consent was obtained from each donor in accordance with the Declaration of Helsinki.

### 2.2. PEVs isolation using tangential flow filtration (TFF)

PEVs were isolated following International Society for Extracellular Vesicles (ISEV) guidelines ^28^. Platelets were divided into two states: (i) resting and (ii) stored. Resting platelets were harvested from PRP immediately after blood collection, while stored platelets were obtained from PRP agitated for 24h at room temperature to mimic platelet storage conditions. PRP was centrifuged at 980 x g for 10 minutes to isolate platelets, which were then resuspended in Tyrode’s buffer. The suspension was centrifuged at 1000 x g for 10 minutes to remove cells, and the resulting platelet-free supernatant was passed through a 0.45 µm membrane filter. PEVs were subsequently isolated by TFF, as previously described by our group^16,27^. In brief, the platelet-free supernatant was processed on a Millipore LabScale TFF system with a Biomax 500 kDa Pellicon filter (Millipore, Billerica, MA). The system was operated under feed pressure <20 psi and retentate pressure <10 psi. Three volume exchanges with calcium-free phosphate-buffered saline (PBS) (500 ml each) were performed before concentrating PEVs into 10 ml PBS.

### 2.3. Quantification of PEV concentration and size distribution by nanoparticle tracking analysis

PEVs size and concentration were measured using a NanoSight NS300 (Malvern Panalytical, UK) equipped with a 488 nm laser and sCMOS camera, as previously described^29^. In brief, samples were diluted 1:100 in 30 kDa-filtered PBS. Particle size measurements were performed at camera level 10 and detection threshold 5. Five consecutive 60s videos were recorded at room temperature while injecting the sample with a syringe pump at flow speed 50. Particle trajectories were analyzed with NTA 2.3 software to determine size distribution and concentration.

### 2.4. Morphological assessment of PEVs by transmission electron microscopy

PEVs were absorbed onto glow-discharged, carbon-coated copper grids (200 mesh; Agar Scientific, UK) for 60 s, dried, and negatively stained with 3% uranyl acetate (System Biosciences, USA) for 60 s. Imaging was performed on a FEI Tecnai 12 TEM at 120 kV with an AMT XR111 CCD camera at the Thomas Jefferson University Core Facility.

### 2.5. Culture conditions of HULEC-5a endothelial cells

Human lung microvascular endothelial cells (HULEC-5a, CRL-3244; ATCC) were cultured in MCDB131 basal medium supplemented with 10% fetal bovine serum (FBS), 1% penicillin – streptomycin, 10 mM glutamine, 10 ng/ml epidermal growth factor, and 1 µg/ml hydrocortisone.

### 2.6. PEVs treatment and measurement of endothelial barrier integrity

Endothelial barrier integrity was evaluated with the xCELLigence live-cell monitoring system, which measures electrical impedance in real time using gold microsensor-coated plates (e-Plate 16). For assay setup, 100 μl of growth medium was added to each well for 10 minutes to establish a background cell index (CI). HULEC-5a cells were then seeded in a plate at a density of 40,000 cells per well and monitored until confluence, indicated by a plateau in the CI curve. Cells were pretreated with PEVs at a concentration of 3.29 x 10^9^ for 60 minutes before stimulation with thrombin (0.2 U/ml). CI values were continuously recorded by the xCELLigence system, normalized to their respective values at the time of thrombin addition, and corrected by subtracting values from untreated controls. The resulting baseline-normalized CI values were used for analysis.

### 2.7. Flow cytometric evaluation of endothelial uptake of PEVs

PEVs were labeled with ExoGlow™-Protein EV Labeling Kit (System Biosciences) at 1:500 dilution for 20 min at 37°C in the dark. After incubation with ExoQuick-TC, samples were centrifuged at 10,000 rpm for 10 min to pellet labeled PEVs. HULEC-5a cells (2 x10^5^ cells/well, 12-well plates) were grown to confluence, then treated with labeled PEVs for 1h. Cells were subsequently stimulated with 0.2 U/ml thrombin for 15 min at 37°C. After washing and trypsinization, cells were fixed in 1% paraformaldehyde with 0.2% BSA and analyzed on a BD LSRFortessa flow cytometer (561 nm laser, PE channel). Data was processed using FlowJo v10.

### 2.8. Western blot analysis of EV surface markers and endothelial junctional proteins

Protein concentration was determined using a BCA assay (Pierce, Thermo Scientific). PEVs (3.29 x 10^9^) were lysed in RIPA buffer containing protease inhibitors, and HULEC-5a cells were seeded in 6-well plates at 1 x 10^5^ cells per well, grown to confluence, pretreated with RP- or SP-PEVs for 1h, and then exposed to 0.2 U/ml thrombin for 1h before lysis in the same buffer. All lysates were centrifuged, and supernatants were stored at −20°C. Proteins from EVs and cells were resolved on 4-20% gradient SDS-PAGE gels, transferred to PVDF membranes, and blocked with 3% BSA. Membranes from EV lysates were probed with antibodies against surface markers (CD9, CD31, CD41, CD63), while those from cell lysates were probed for junctional proteins (ZO-1, VE-cadherin, occludin, JAM-A, claudin-5). GAPDH was used as a loading control. HRP-conjugated secondary antibodies were applied; bands were visualized with ECL substrate (Thermo Fisher), imaged on a Chemidoc system (Bio-Rad), and quantified by densitometry using ImageJ.

### 2.9. Visualization of endothelial junctional protein localization

HULEC-5a cells were cultured on 0.1% gelatin-coated coverslips until confluent. Cells were pretreated with PEVs for 1h, followed by 0.2 U/ml thrombin challenge for 1h. After fixation in 4% paraformaldehyde, cells were permeabilized with 0.25% Triton X-100 and blocked with 2% BSA. Coverslips were incubated overnight at 4°C with primary antibodies against junction proteins (ZO-1, VE-cadherin, occludin, JAM-A, claudin-5). After PBS washes, Alexa Fluor 555-conjugated secondary antibodies were applied for 2h at room temperature. Nuclei were counterstained with DAPI mounting medium. Images were acquired with an EVOS FL inverted microscope (60X).

### 2.10. Detection of intracellular reactive oxygen species levels in endothelial cells

ROS levels were measured using 2′,7′-dichlorodihydrofluorescein diacetate (H2DCFDA), a cell-permeable probe that is oxidized by intracellular ROS to form the fluorescent compound 2′,7′-dichlorofluorescein (DCF). To assess ROS production, HULEC-5a cells were pretreated with RP-PEVs or SP-PEVs, exposed to thrombin for 1h, then harvested, resuspended in PBS, and incubated with 10 µg/ml H2DCFDA for 30 min at 37 °C in the dark. Fluorescence was measured on a BD Accuri C6 Plus flow cytometer (492 nm excitation, 517 nm emission), and mean fluorescence intensity (MFI) was used as the readout of ROS levels.

### 2.11. Gene expression analysis of endothelial inflammatory mediators by qRT-PCR

HULEC-5a cells were seeded at 1 x 10^5^ cells per well in 12-well plates, treated with PEVs, and then exposed to 0.2 U/ml thrombin for 1h. Total RNA was extracted using the RNeasy Mini Kit (Qiagen), and yield and purity were determined with a Nanodrop spectrophotometer based on the 260/280 ratio. One microgram of RNA was reverse transcribed into cDNA with the iScript cDNA kit (Bio-Rad). Quantitative PCR was carried out on a QuantStudio 3 system using SYBR™ Green Master Mix (Thermo Fisher) and gene-specific primers (**Table 1**), with GAPDH serving as the internal control. Cycling conditions were 50°C for 2 min, 95°C for 10 min, and 45 cycles of 95°C for 15s and 50°C for 1 min. Relative expression was calculated using the 2−ΔΔCt method. Analyses were performed on sex-specific pools under resting and stored conditions, and results are presented as a heat map showing mean fold change relative to untreated samples.

**Table 1.**
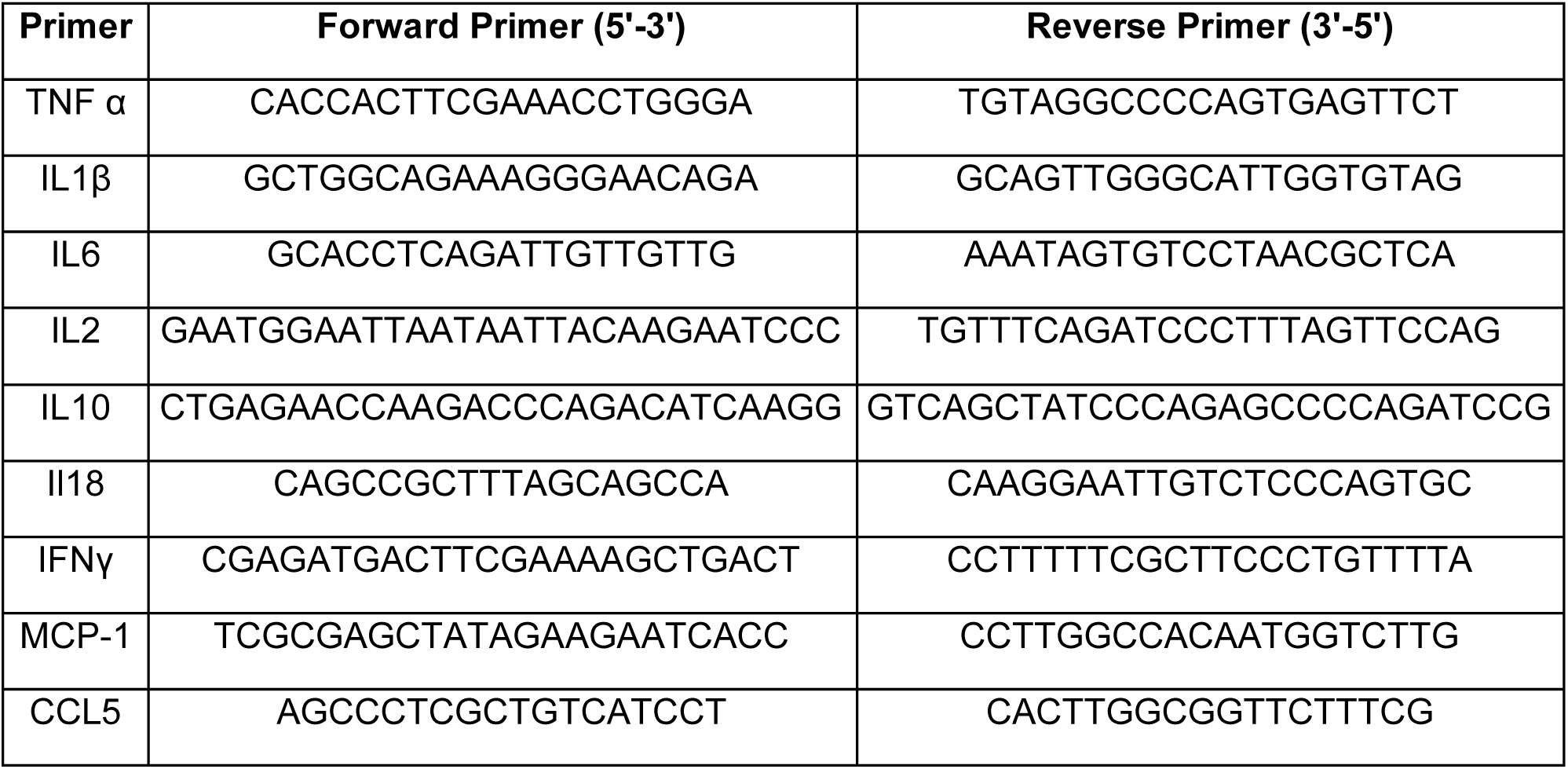
Primer sequences for qRT-PCR analysis of endothelial inflammatory mediators.

### 2.12. Data analysis and statistical methods

Comparisons between groups were determined by one-way ANOVA. A p-value ≤ 0.05 was considered statistically significant. Analyses were performed using GraphPad Prism 9 (GraphPad, San Diego, CA). All experiments were independently repeated at least three times. Data are expressed as mean ± standard error of the mean (SEM).

## 3. Results

### 3.1. PEVs display consistent morphology with sex-specific molecular variation

TEM analysis showed intact membranes and electron-dense interiors, predominantly 100-200 nm in diameter in both RP-PEVs and SP-PEVs samples from male and female donors, with no systematic morphological differences observed between conditions (**Fig. 1A**). NTA confirmed comparable size distributions across donors but revealed that average size of SP-PEVs from female donors were significantly smaller compared to RP-PEVS (137.7 ± 44.9 nm vs. 190.5 ± 20.9 nm, p=0.04). No difference in size was detected between RP- and SP-PEVs in male donors (151.9 ± 55.1 nm vs. 156.3 ± 44.2 nm) (**Fig. 1B**). Quantitatively, 70% of PEVs in the male group were 0-150 nm in size, compared with 50% in the female group. In contrast, 30% of PEVs in the female group measured 150-200 nm, whereas only 10% fell into this range in the male group, indicating that PEVs from females tend to be relatively larger. A detailed summary of platelet and PEV characteristics across donors is presented in **Table 2**. Western blot analysis confirmed the expression of platelet source markers (CD41, CD31) as reported previously^30^ and EV markers (CD9, CD63) in all samples. CD41 and CD31 were consistently detected in PEVs from all donors, although band intensities varied. Notably, CD63 expression was higher in male RP-PEVs than in female RP-PEVs, suggesting sex-dependent regulation. In contrast, CD9 expression was lower in female RP-PEVs compared with SP-PEVs, indicating storage-related variability. No significant differences in CD9 expression were observed between male RP-PEVs and SP-PEVs (**Fig. 1C**).

**Figure 1.**
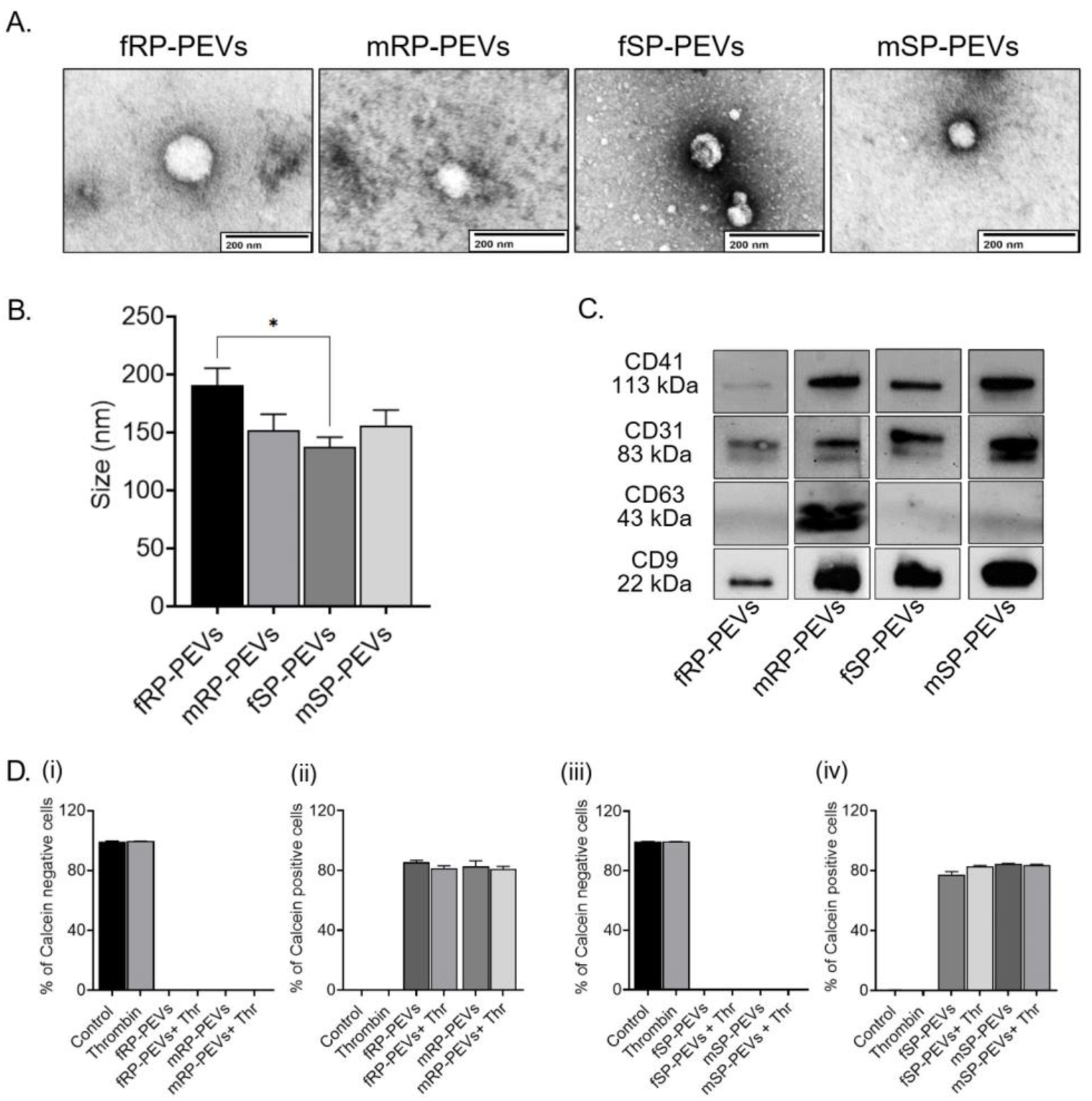
Biophysical and molecular characterization of RP- and SP-PEVs. **(A)** TEM images of isolated RP- and SP-PEVs, revealing round morphology with diameters of 100-400 nm. Scale bars, 200 nm. **(B)** NTA showing size distribution and concentration of PEVs. Significant difference in size distribution observed between resting and stored female donors. **(C)** Western blot detection of canonical EV markers (CD9, CD63), platelet marker (CD41), and platelet-endothelial marker (CD31) in PEV samples. **(D)** PEVs uptake by endothelial cells is not influenced by donor sex. HULEC-5a cells were treated with fluorescently labeled RP- or SP-PEVs from male and female donors, followed by thrombin stimulation. Uptake was quantified via flow cytometry. Quantification of PEV-positive cells shows no significant sex-based differences in uptake across groups with RP-**(i-ii)**- or SP-PEVs **(iii-iv)**. Data represents mean ± SEM from triplicate experiments. *p < 0.05. MW, molecular weight; f, female; m, male. resting (fRP-PEVs, mRP-PEVs) and stored (fSP-PEVs, mSP-PEVs).

**Table 2.** Donors, platelet count, and corresponding PEVs information.

| Donor | Sex | State | Platelet count/μl | PEVs/ml | Size (in nm) |
| --- | --- | --- | --- | --- | --- |
| 1 | Female | Resting | 4,86,0000 | 1.54e + 009 | 221.4 ± 12.0 |
| 2 | Female | Resting | 6,700,000 | 7.9e + 010 | 136.9 ± 55.5 |
| 3 | Female | Resting | 1,140,000 | 1.99e + 009 | 212.1 ± 8.50 |
| 4 | Female | Resting | 4,025,00 | 4.90e + 009 | 188.0 ± 8.30 |
| 5 | Female | Resting | 3,815,000 | 5.51e + 010 | 194.9 ± 20.2 |
| 6 | Female | Stored | 2,480,000 | 1.61e + 008 | 156.8 ± 7.40 |
| 7 | Female | Stored | 6,300,000 | 5.9e + 010 | 137.3 ± 59.7 |
| 8 | Female | Stored | 1,560,0000 | 9.9e + 010 | 141.9 ± 59.3 |
| 9 | Female | Stored | 6,000,000 | 1.4e + 011 | 144.8 ± 63.3 |
| 10 | Female | Stored | 4,130,000 | 1.7e + 009 | 107.9 ± 34.8 |
| 11 | Male | Resting | 3,815,000 | 4.5e + 008 | 206.6 ± 36.7 |
| 12 | Male | Resting | 7,500,000 | 9.3e + 010 | 141.3 ± 55.3 |
| 13 | Male | Resting | 4,300,000 | 3.0e + 009 | 129.0 ± 62.0 |
| 14 | Male | Resting | 6,200,000 | 1.2e + 011 | 140.4 ± 58.6 |
| 15 | Male | Resting | 6,700,000 | 7.8e + 010 | 142.2 ± 62.9 |
| 16 | Male | Stored | 8,000,000 | 2.85e + 010 | 204.7 ± 6.90 |
| 17 | Male | Stored | 2,018,000 | 3.27e + 009 | 135.6 ± 12.5 |
| 18 | Male | Stored | 5,350,000 | 5.4e + 010 | 147.6 ± 67.4 |
| 19 | Male | Stored | 6,950,000 | 2.8e + 010 | 159.4 ± 77.0 |
| 20 | Male | Stored | 4,695,000 | 2.2e + 010 | 134.4 ± 57.2 |

\* PEVs size – average ± standard error of mean

### 3.2. PEVs are efficiently internalized by endothelial cells independent of donor sex or platelet storage conditions

To evaluate endothelial interaction and internalization, we quantified uptake of labeled PEVs in HULEC-5a cells by flow cytometry. Labeled PEVs treatment shifted cell populations to the positive gate, confirming uptake. The proportion of PEV-positive cells exceeded 80% and did not change after thrombin treatment. Uptake was comparable between RP- and SP-PEVs and did not differ by donor sex (**Fig. 1D**).

### 3.3. Female PEVs provide superior protection against thrombin-induced endothelial barrier disruption

RP-PEVs markedly preserved endothelial barrier integrity following thrombin challenge across all donors (p < 0.001; **Fig. 2A**). Both female and male RP-PEVs restored normalized CI to near-baseline levels, demonstrating strong and consistent protection independent of donor variability. In contrast, SP-PEVs provided less protection overall. Female SP-PEVs consistently maintained endothelial barrier integrity across all donors, improving CI recovery after thrombin exposure. Male SP-PEVs demonstrated inconsistent protection, with barrier integrity preserved in only four of five donor preparations (**Fig. 2B**). These results indicate that RP-PEVs are more effective than SP-PEVs in mitigating thrombin-induced endothelial disruption. Sex-stratified analysis confirmed these trends (**Fig. 2C**). Female RP-PEVs conferred the greatest protection, enhancing TEER recovery by >99% compared with 88.24% for male RP-PEVs. Female SP-PEVs improved recovery by 91.76%, whereas male SP-PEVs achieved only 56.47%, showing no significant difference from thrombin treatment alone. Treatment of unchallenged endothelial monolayers with either RP-PEVs or SP-PEVs did not affect baseline CI, confirming that the vesicles themselves did not compromise barrier integrity. Individual CI data for all donors are presented in **Table 3**. These findings demonstrate that RP-PEVs provide superior protection against thrombin-induced endothelial dysfunction compared with SP-PEVs. The effect is further modulated by donor sex, with female RP-PEVs showing the most potent and consistent preservation of endothelial barrier function.

**Figure 2.**
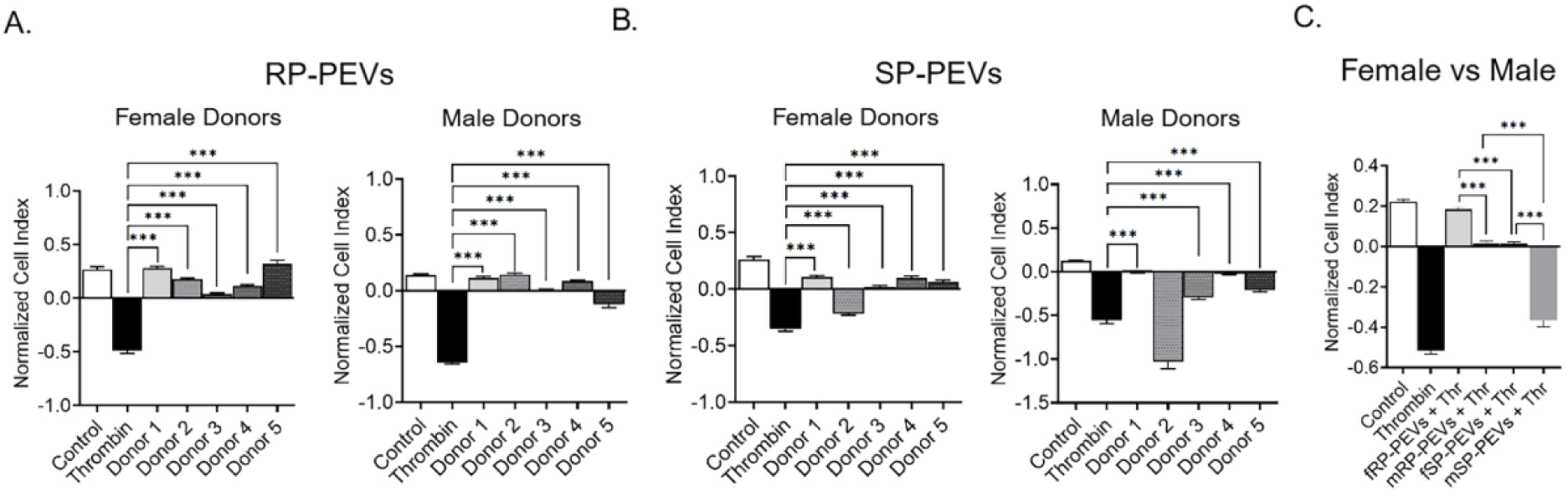
Sex- and platelets storage-dependent variation in PEV-mediated preservation of endothelial barrier integrity. HULEC-5a monolayers were treated with PEVs prior to thrombin challenge. TEER was monitored. **(A-B)** Quantification of TEER responses stratified by donor for cells treated with RP-**(A)** or SP-PEVs **(B)** from female and male donors. **(C)** Pooled data showing cumulative barrier-protective effects of RP-PEVs and SP-PEVs. Data are shown as mean ± SEM. *p < 0.05, **p < 0.01, ***p < 0.001, ****p < 0.0001, ns = not significant.

**Table 3.** Endothelial cell impedance expressed as cell index (CI).

| **State** | **Donor 1** | **Donor 2** | **Donor 3** | **Donor 4** | **Donor 5** |
| --- | --- | --- | --- | --- | --- |
| Female<br>(Resting) | 0.27 ± 0.17 | 0.17 ± 0.11 | 0.03 ± 0.09 | 0.11 ± 0.10 | 0.32 ± 0.29 |
| Male<br>(Resting) | 0.05 ± 0.04 | 0.19 ± 0.08 | 0.02 ± 0.02 | 0.11 ± 0.05 | -0.30 ± 0.21 |
| Female<br>(Stored) | 0.10 ± 0.13 | -0.21 ± 0.09 | 0.02 ± 0.10 | 0.10 ± 0.13 | 0.06 ± 0.13 |
| Male<br>(Stored) | 0.01 ± 0.05 | -1.24 ± 0.64 | -0.35 ± 0.18 | -0.01 ± 0.05 | -0.24 ± 0.13 |

### 3.4. PEVs restore endothelial junctional protein expression and localization in a manner dependent on donor sex and platelet storage conditions

We next assessed whether PEVs preserve endothelial junctional integrity by examining the expression and localization of tight and adherens junction proteins. Western blot analysis quantified ZO-1, VE-cadherin, occludin, JAM-A, and claudin-5 in thrombin-stimulated endothelial cells pretreated with PEVs under fresh (**Fig. 3A)** and stored **(Fig 4A**) conditions. Band intensities were normalized to GAPDH and expressed as fold change relative to control (set to 1). Thrombin exposure markedly reduced the expression of all junctional proteins, confirming disruption of endothelial barrier components. Pretreatment with PEVs attenuated these losses in a sex-dependent manner with female PEVs produced stronger and more consistent restoration of junctional protein expression than male PEVs. Among RP-PEVs, female RP-PEVs substantially recovered VE-cadherin (1.10 ± 0.15-fold), and JAM-A (0.92 ± 0.30-fold), with moderate improvement in ZO-1 (0.52 ± 0.06-fold), occludin (0.64 ± 0.06-fold) and claudin-5 levels (0.50 ± 0.01-fold). Male RP-PEVs also improved ZO-1 (0.65 ± 0.06-fold), VE-cadherin (0.73 ± 0.14-fold) and occludin (0.75 ± 0.14-fold) but had minimal effects on the remaining junctional markers, indicating partial and less coordinated protection. A similar trend was observed for SP-PEVs: female SP-PEVs provided the most comprehensive recovery across all proteins, markedly increasing ZO-1 (0.96 ± 0.18-fold), VE-cadherin (1.36 ± 0.08-fold), occludin (1.08 ± 0.11-fold), JAM-A (0.79 ± 0.02-fold), and claudin-5 (0.66 ± 0.03-fold). In contrast, male SP-PEVs selectively increased claudin-5 (0.95 ± 0.04-fold) but partially restored ZO-1, VE-cadherin, or JAM-A, resulting in incomplete junctional recovery. Fold-change values for all markers are summarized in **Table 4**.

**Figure 3.**
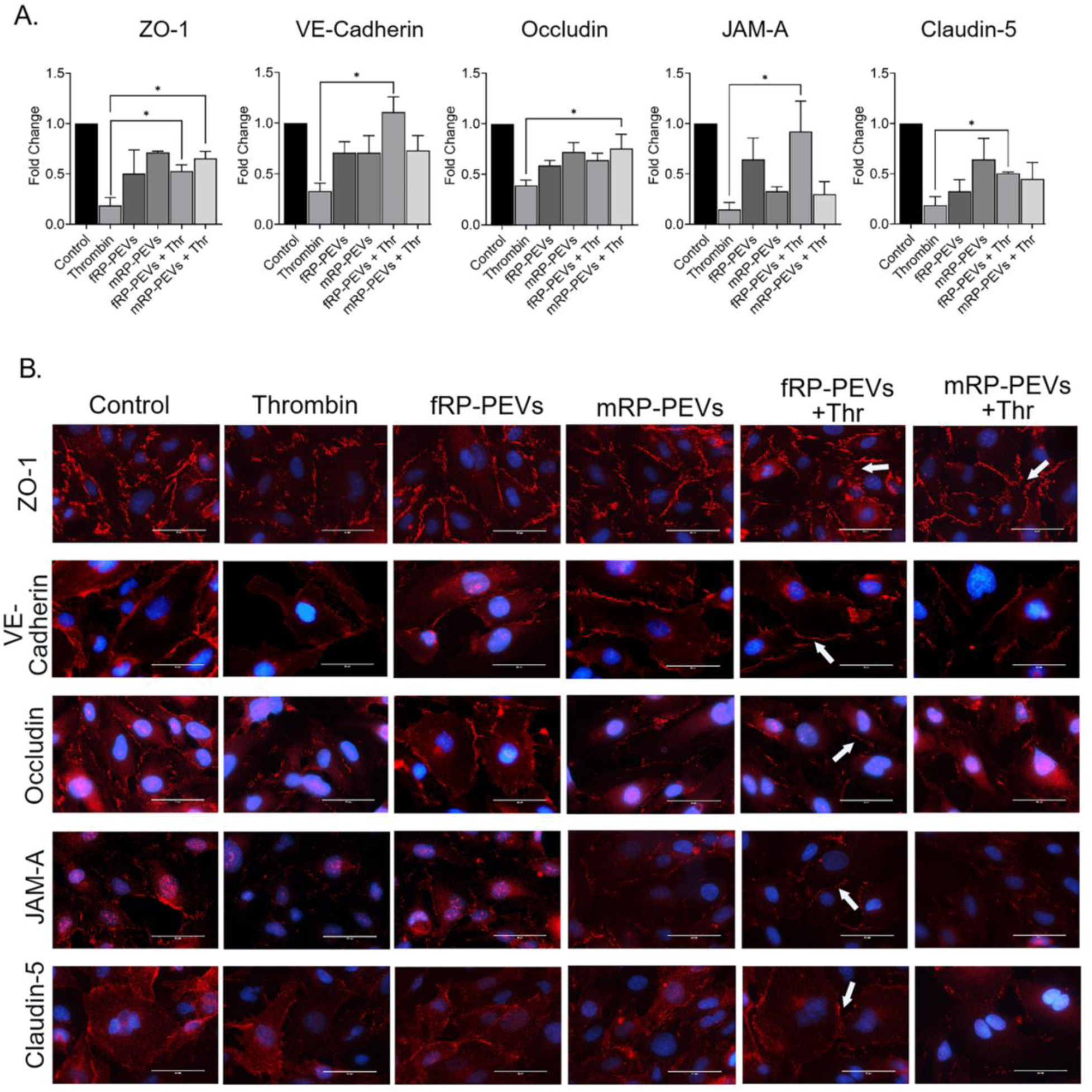
Effects of RP-PEVs on endothelial junction integrity in thrombin-challenged cells. **(A)** HULEC-5a cells after treatment with RP-PEVs. GAPDH was used as a loading control. Densitometric analysis shows fold changes relative to control. PEVs, particularly from female donors, preserved junctional protein levels following thrombin exposure. Data are shown as mean ± SEM *p < 0.05, **p < 0.01, ns = not significant. **(B)** Immunofluorescent staining for junction proteins (red) in HULEC-5a cells treated with RP-PEVs and thrombin. Female PEVs preserved the junctional integrity. Junctional localization indicated by white arrows. Scale bars, 50 μm. Data represent mean ± SEM. *p < 0.05, **p < 0.01 vs. control.

**Figure 4.**
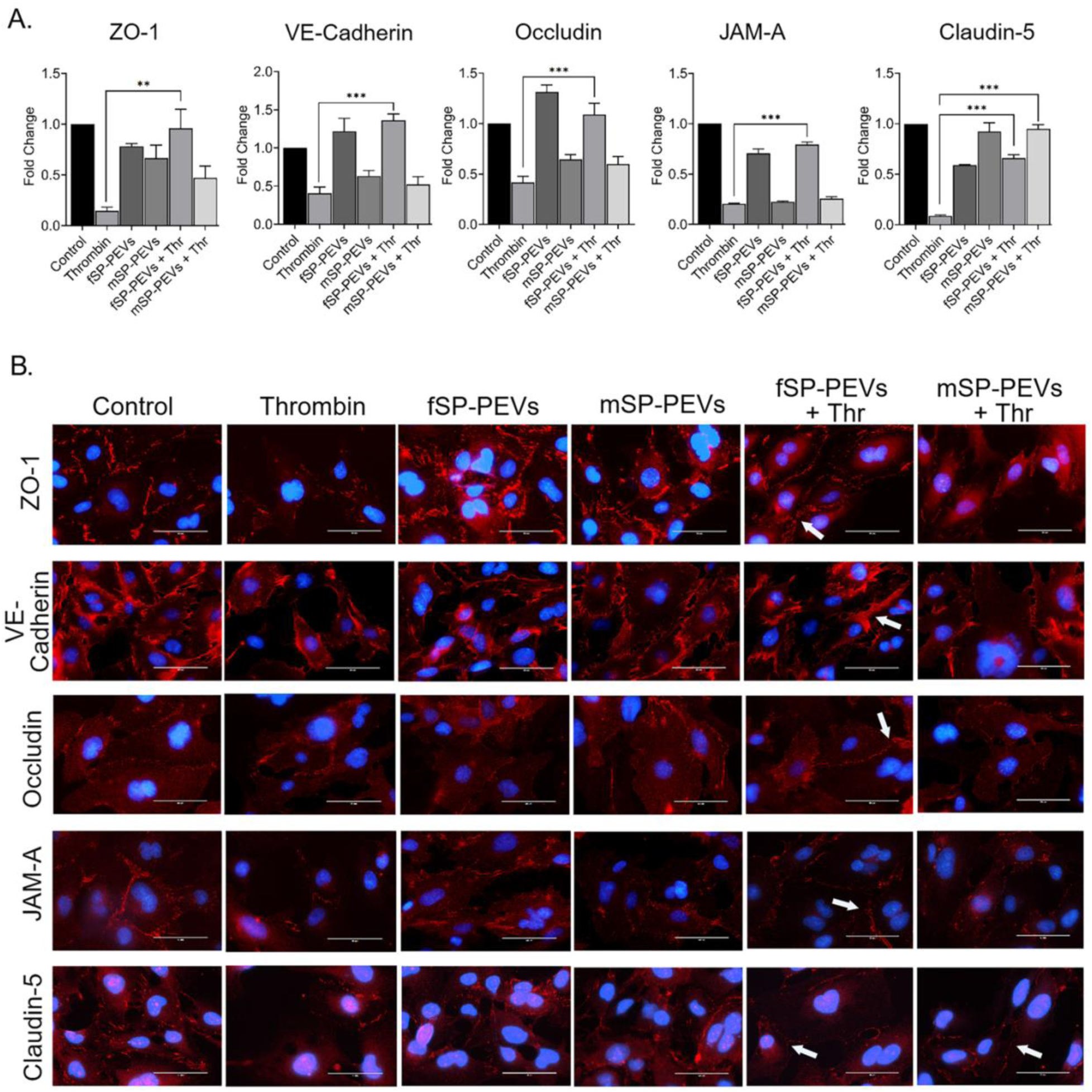
Effects of SP-PEVs on endothelial junction integrity in thrombin-challenged cells. **(A)** HULEC-5a cells after treatment with RP-PEVs. GAPDH was used as a loading control. Densitometric analysis shows fold changes relative to control. Female-derived PEVs preserved junctional protein levels than male PEVs. Data are shown as mean ± SEM *p < 0.05, **p < 0.01, ns = not significant. **(B)** Immunofluorescent staining for junction proteins (red) in HULEC-5a cells treated with SP-PEVs and thrombin. Effects of female SP-PEVs were pronounced than male donors. Junctions indicated by white arrows. Scale bars, 50 μm. Data represent mean ± SEM. *p < 0.05, **p < 0.01 vs. control.

**Table 4.** Protein expression changes in junctional proteins determined with western blot analysis.

| **State** | **ZO-1** | **VE-Cadherin** | **Occludin** | **JAM-A** | **Claudin-5** |
| --- | --- | --- | --- | --- | --- |
| Female<br>(Resting) | 0.52 ± 0.06 | 1.10 ± 0.15 | 0.64 ± 0.06 | 0.92 ± 0.30 | 0.50 ± 0.01 |
| Male<br>(Resting) | 0.65 ± 0.06 | 0.73 ± 0.14 | 0.75 ± 0.14 | 0.29 ± 0.12 | 0.44 ± 0.16 |
| Female<br>(Stored) | 0.96 ± 0.18 | 1.36 ± 0.08 | 1.08 ± 0.11 | 0.79 ± 0.02 | 0.66 ± 0.03 |
| Male<br>(Stored) | 0.47 ± 0.11 | 0.52 ± 0.10 | 0.59 ± 0.67 | 0.25 ± 0.01 | 0.95 ± 0.04 |

\* *Fold changes compared to control (fold change ± standard error of mean)*

Immunocytochemical analysis confirmed the biochemical findings and showed clear sex- and storage-dependent differences in endothelial junction organization after treatment with fresh (**Fig. 3B) and** stored **(Fig 4B**) PEVs. In untreated endothelial monolayers, ZO-1 and VE-cadherin showed continuous, sharply defined borders typical of intact tight and adherens junctions. Occludin, JAM-A, and claudin-5 were evenly distributed along cell-cell contacts, indicating a cohesive and organized junctional network. Thrombin treatment caused marked disruption. ZO-1 and VE-cadherin staining became fragmented and discontinuous, and occludin, JAM-A, and claudin-5 lost their junctional localization and appeared as diffuse or cytoplasmic signals, consistent with junction disassembly and loss of barrier integrity. Pretreatment with PEVs reduced these thrombin-induced changes, and the extent of protection varied with donor sex and platelet storage condition. Endothelial cells treated with female RP-PEVs closely resembled control cells (**Fig 3B**). ZO-1 and VE-cadherin borders remained near-continuous and well defined, and occludin, JAM-A, and claudin-5 were relocalized to the cell borders, showing some restoration of junctional structure. Male RP-PEVs provided partial recovery showing fragmented ZO-1 and VE-cadherin borders. Occludin and JAM-A signals remained mainly cytoplasmic. A similar trend was seen with PEVs from stored platelets (**Fig 3B**). Female SP-PEVs preserved junctional organization after thrombin exposure. Cells pretreated with these vesicles showed continuous ZO-1 and VE-cadherin staining (**Fig 4B**). Partial recovery of occludin, claudin-5 and JAM-A at cell contacts was observed. Male SP-PEVs were less protective with irregular borders for ZO-1 and VE-cadherin, while other junctional proteins remained discontinuous and cytoplasmic (**Fig 4B**).

### 3.5. Female RP-PEVs most effectively suppress endothelial ROS accumulation

The increased fluorescence intensity in thrombin-treated cells indicates elevated ROS levels, which were effectively lowered upon PEVs pretreatment in a sex- and state-dependent manner (**Fig. 5**). PEVs pretreatment significantly suppressed ROS generation compared to the thrombin. Female-derived RP- and SP-PEVs suppressed ROS more effectively than male PEVs (p < 0.01). Relative to thrombin treatment, ROS level decreased by female RP-PEVs (**Fig. 5A**) by 38.84% versus 26.81% after female SP-PEVs treatment (**Fig. 5B**). Male donors from RP- and SP-PEV groups showed no significant reductions (**Fig. 5A-B**). These findings indicate that PEV modulation of oxidative stress depends on donor sex and platelet storage, with RP-female PEVs providing superior protection against ROS accumulation among all the groups.

**Figure 5.**
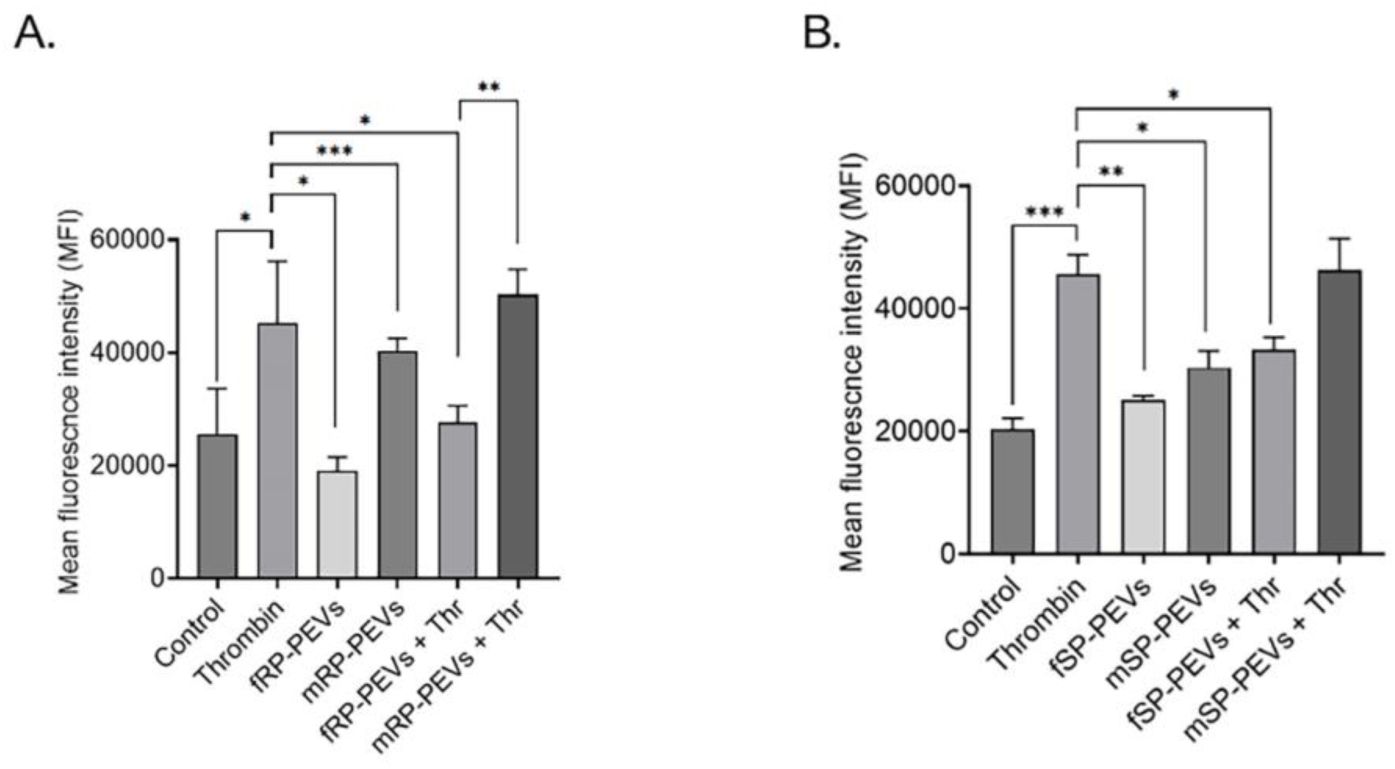
Effects of PEVs in attenuationg thrombin-induced ROS generation in endothelial cells. ROS production in HULEC-5a cells was quantified using H2DCFDA after pretreatment with PEVs and thrombin exposure. Comparison of pooled RP-**(A)** vs. SP-**(B)** PEVs. Data represent mean ± SEM from three independent experiments. *p < 0.05, **p < 0.01, ***p < 0.001. f, female; m, male.

### 3.6. PEVs suppress thrombin-induced endothelial inflammatory mediators with donor sex influencing specific pathways

To assess the effect PEVs on thrombin-induced endothelial inflammatory mediators, we measured mRNA levels of pro-inflammatory cytokines and chemokines in endothelial cells pretreated with PEVs and then challenged with thrombin. The heat map displays the average fold change compared to control **(Fig. 6)**. TNF-α was significantly downregulated by PEVs regardless of sex of the donors. Male RP-PEVs and SP-PEVs produced the strongest reduction. IL-1β, which increases permeability via VEGF induction, was reduced after PEV pretreatment, with female SP-PEVs showing the greatest effect (0.94 ± 0.08). IL-2, a cytokine that disrupts adherens junctions and increases permeability^31^, was reduced across all PEV groups, specifically male RP-PEVs group, indicating broad protection of adherens junctions. IL-10, which inhibits TNF-α, IFN-γ, and IL-6^32^, increased significantly with both RP-(p < 0.05) and SP-(p < 0.001) PEVs; the largest increase occurred with male RP-PEVs (2.69 ± 0.88). IL-18, a mediator linked to vascular injury, was significantly downregulated by male RP-PEVs (p < 0.0001). IFN-γ, a potent disruptor of barrier integrity, was also reduced, with male RP-PEVs, male and female SP-PEVs. MCP-1 was suppressed by PEVs, with greater reduction in the male RP-PEVs group (0.55 ± 0.09 vs. thrombin 2.29 ± 0.44). CCL5 was not significantly altered across most individual donors. Lastly, the male RP-PEVs efficiently lowered IL-6 expression. Cumulatively, these findings summarize that male donors under resting conditions were effective in mitigating inflammation. Collectively, male RP-PEVs showed higher proficiency among all other groups.

**Figure 6.**
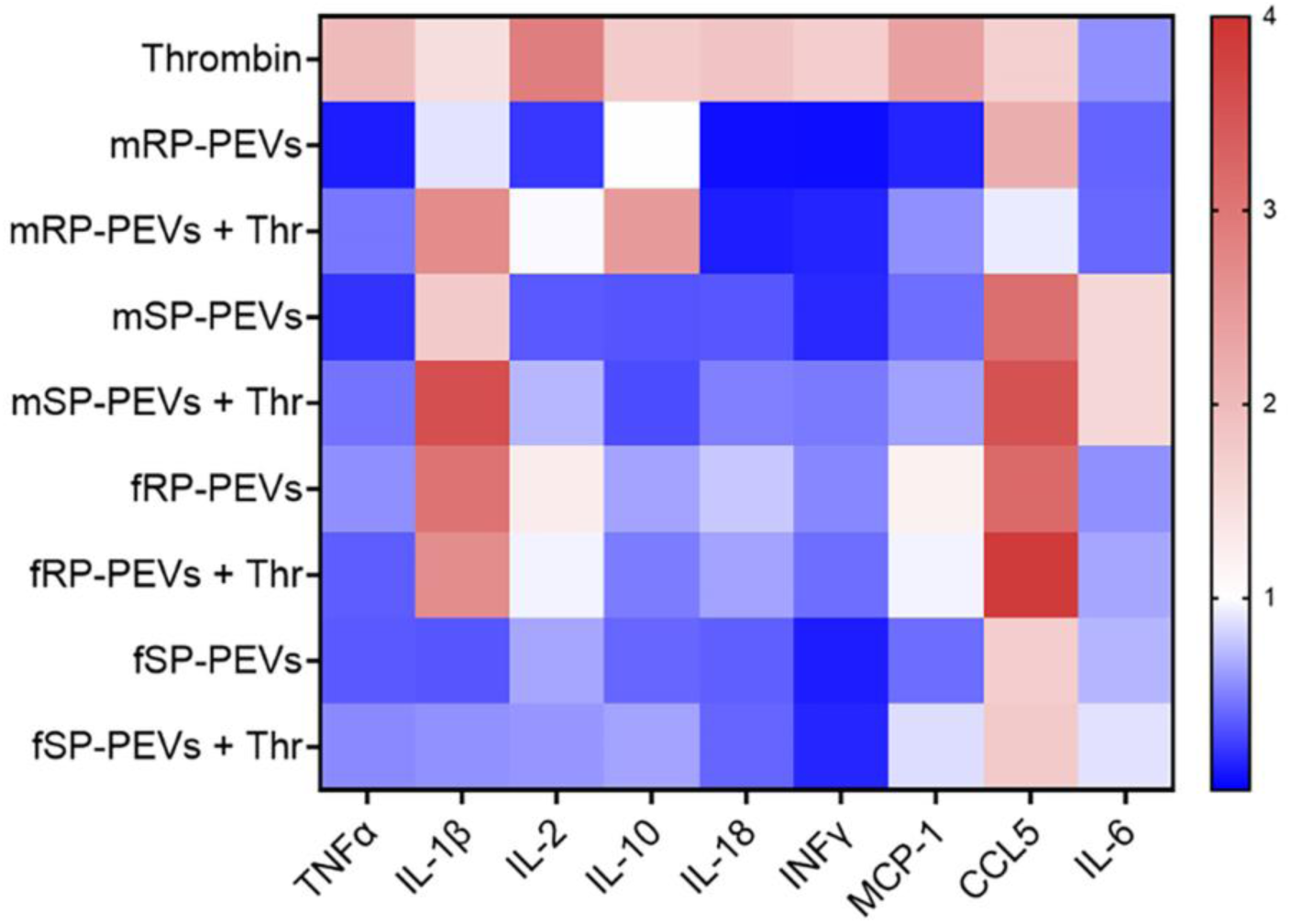
PEVs differentially modulate endothelial inflammatory mediator expression. Heat map summarizing quantitative RT-PCR mRNA expression of inflammatory mediators (TNF-α, IL-1β, IL-6, IL-2, IL-10, IL-18, IFN-γ, MCP-1, and CCL5) in HULEC-5a cells treated with RP-g or SP-PEVs from male and female donors, followed by thrombin stimulation. Expression levels were normalized to GAPDH and calculated using the 2^−ΔΔCt method. Data are presented as fold-change relative to control (mean ± SEM). Color scale denotes fold-change values, shown at right. Notable findings include reduced TNF-α, IL-2, IL-18, IFN-γ, and MCP-1 expression with male RP-PEVs, and modulation of IL-10 by the male RP-PEV group.

## 4. Discussion

PEVs are biologically active mediators that influence vascular repair^4^. This study demonstrates that PEV bioactivity depends on both donor sex and platelet storage state with RP-PEVs from female donors producing the most consistent endothelial-protective phenotype *in vitro*. In human lung endothelial cells challenged with thrombin, RP-PEVs treatment preserved TEER markedly and female RP-PEVs yielded the highest median TEER recovery, accompanied by restoration of key junctional proteins and substantial suppression of thrombin-induced ROS (**Figs.2-5**). These convergent functional readouts argue that female RP-PEVs deliver a coordinated set of bioactive cargos that promote junctional reassembly and redox homeostasis, producing functionally meaningful barrier protection in this model.

The mechanistic basis for the sex-specific disparity is likely multifactorial. Estrogen influences platelet biology and vascular tone^33^, and female platelets release vesicles enriched in miRNAs such as miR-223 and miR-126^34–39^. These miRNAs regulate endothelial activation, stabilize junctional complexes, and promote cytoskeletal remodeling^40^. Additionally, sex-related differences in platelet mitochondrial function may also contribute. Female platelets exhibit higher oxidative phosphorylation rates, greater mitochondrial reserve capacity, and enhanced resistance to oxidative stress compared with male platelets^41,42^. These metabolic features could influence the composition and functional activity of released vesicles, predisposing female PEVs to carry antioxidant and junction-stabilizing cargo, consistent with our observations of reduced ROS, improved junctional protein localization, and suppression of inflammatory cytokine.

We observed that despite equivalent uptake of ExoGlow-labeled PEVs across donor sex and platelet storage conditions, their downstream functional effects differed substantially. These findings indicate that therapeutic efficacy depends on vesicle cargo composition rather than overall uptake efficiency. Previous studies have reported similar observations, showing that comparable internalization rates can lead to distinct biological responses depending on the RNA or protein content of extracellular vesicles^43,44^. This distinction underscores that measuring uptake alone is insufficient to infer bioactivity; functional potency reflects the quality and composition of transferred cargo rather than the quantity of vesicles internalized.

We also evaluated the consequences of platelet storage using a 24-hour rocking protocol to mimic mechanical stimulation during blood bank storage. PEVs generated under these conditions exhibited reduced therapeutic activity, reflected by weaker barrier preservation, diminished junctional protein expression, and impaired suppression of ROS and cytokine release. Proteomic studies suggest that proteins such as BICD1, a dynein adaptor regulating Golgi–ER transport and PAR1 internalization, and flotillin, a scaffolding protein involved in membrane organization and signaling, are enriched in resting platelet vesicles^45,46^. These observations support the notion that quiescent PEVs retain trafficking and structural elements enhancing endothelial protection. In contrast, prolonged platelet storage alters vesicle cargo by promoting the accumulation of pro-inflammatory and pro-thrombotic mediators within EVs⁴⁶, producing a more inflammatory secretome^47^. Changes in vesicle cargo that occur during platelets storage explain why PEVs obtained from stimulated platelets in this study provide less protection. This highlights the importance of consistent bioprocessing and handling, since even slight differences in how vesicles prepared can affect their function.

Interestingly, female PEVs were less affected by storage-induced loss of efficacy, suggesting a sex-specific resilience to stress-induced vesicle reprogramming. It is possibly due to estrogen-mediated stabilization of vesicle biogenesis pathways or reduced incorporation of harmful cargo^42^. This aligns with studies showing that estrogen enhances platelet antioxidant defenses and alters granule content, favoring the release of anti-inflammatory and anti-apoptotic factors^48,49^.

Restoration of endothelial junctions is a critical endpoint in vascular repair^50^. VE-cadherin, ZO-1, occludin, JAM-A, and claudin-5 maintain intercellular cohesion, and their thrombin-induced loss in our study reflects RhoA/ROCK-mediated cytoskeletal contraction and junctional disassembly^51,52^. Pretreatment with PEVs mitigated this disruption, restoring both expression and localization of junctional proteins, with efficacy dependent on donor sex and platelet storage. Female RP-PEVs achieved the most complete recovery. They restored VE-cadherin and ZO-1 continuity and relocalized occludin, JAM-A, and claudin-5 to cell-cell borders, resembling untreated controls. This coordinated restoration indicates preservation of junctional integrity rather than selective protein upregulation. VE-cadherin and ZO-1 form the structural backbone of adherens and tight junctions, respectively, and their coordinated expression is essential for barrier reassembly^53,54^. Recovery of these markers after PEVs treatment suggests stabilization of intercellular contacts and reversal of thrombin-induced cytoskeletal tension. Sex differences in PEVs efficacy likely reflect biological distinctions in platelet composition and metabolism. Female platelets have greater mitochondrial reserve and higher antioxidant activity^34,42^, which may influence vesicle cargo. Proteomic analyses of PEVs show enrichment of cytoskeletal, metabolic, and adhesion-regulating proteins, including integrins and actin-binding components, which modulate junction organization^55^. Enhanced abundance or stability of these cargoes in female PEVs underlie more robust junctional restoration. Coordinated VE-cadherin and ZO-1 localization further suggests PEV-mediated transfer of regulatory molecules that sustain adherens–tight junction crosstalk^51,56^. Reduced efficacy of PEVs from stored platelets aligns with known storage effects, including oxidative stress, degranulation, and cytoskeletal remodeling, which alter EV protein and miRNA content^57,58^. These changes likely diminish adhesion-supportive and antioxidant cargo, explaining attenuated protection by stored PEVs. Nonetheless, stored female PEVs retained partial restorative capacity, reflecting greater resilience of female platelets to storage-induced activation, consistent with reports of sex-dependent platelet metabolism and oxidative balance^49^.

Oxidative stress is both a driver and a result of endothelial dysfunction^59^. Thrombin and other permeability-inducing stimuli activate NADPH oxidase and mitochondrial ROS production, causing cytoskeletal collapse and junctional disruption through the RhoA/ROCK signaling pathway^60^. Our findings show that female RP-PEVs significantly reduced ROS, suggesting delivery of antioxidant cargo. Platelet vesicles are known to carry enzymes such as peroxiredoxins, thioredoxin, and glutathione-related enzymes^61,62^. Differences in platelet mitochondrial metabolism and estrogen-regulated antioxidant pathways^34,49^ can influence the enrichment of these factors in vesicles, potentially explaining the sex- and storage-dependent effects observed. In addition to transferring enzymatic antioxidants, PEVs may also influence endothelial metabolism or redox-sensitive transcription factors^63^, thereby enhancing endogenous endothelial defense responses.

Thrombin induces endothelial inflammation by upregulating pro-inflammatory cytokines and chemokines, which compromise vascular integrity^55^. In our study, pretreatment with PEVs attenuated this inflammatory response, with donor sex influencing the efficacy and specificity of cytokine modulation^34^. TNF-α, a central mediator of endothelial activation, was significantly downregulated by male RP-PEVs. This reduction likely contributes to enhanced stabilization of cell-cell junctions, reducing endothelial permeability^64^. Reduction of TNF-α, along with IL-1β (lowered expression in female SP-PEVs), promotes the expression of adhesion molecules^65^. Male RP-PEVs also enhanced the expression of anti-inflammatory IL-10, suggesting the suppression of other inflammatory markers^66^. IL-2, IL-18, and IFN-γ were also suppressed, predominantly by male RP-PEVs, in conjunction with MCP-1 and IL-6, highlighting coordinated modulation of both chemotactic and cytokine-driven inflammatory pathways^31^. These findings indicate that male RP-PEVs are particularly effective at mitigating thrombin-driven endothelial inflammation, while female and stored PEVs exhibit selective, pathway-specific effects.

Our study challenges the assumption that EVs are functionally interchangeable. Donor characteristics and platelet storage generate meaningful heterogeneity that directly impacts therapeutic consistency and efficacy. These findings underscore the need for functional potency assays, analogous to platelet aggregometry or coagulation testing, rather than relying solely on vesicle size, concentration, or marker expression. Functional readouts, including barrier restoration, ROS suppression, and cytokine modulation, should be incorporated into manufacturing and quality control pipelines. These insights also open opportunities for rational design of engineered PEVs, selectively enriched with protective cargo or mimetic molecules to replicate the beneficial molecular signatures of female RP-PEVs, advancing precision nanomedicine approaches.

A limitation of this study is the absence of detailed molecular profiling of PEVs. High-throughput proteomic, transcriptomic, and lipidomic analyses will be required to identify predictive biomarkers of efficacy. Nevertheless, these results establish a foundation for precision-based PEVs therapies, emphasizing donor selection and standardized bioprocessing to enhance therapeutic consistency.

In conclusion, PEVs are not functionally interchangeable but reflect donor- and platelets storage-dependent heterogeneity with direct implications for therapeutic efficacy. Female PEVs consistently demonstrated superior endothelial protection, highlighting the importance of sex-aware donor selection and production strategies.

## Acknowledgments

This work was supported by the U.S. Department of Defense grants W81XWH2110682 (to Amit K. Srivastava) and W81XWH2110683 (to Jessica C. Cardenas).

